# TurboID proximity interactome mapping reveals NR2E3 association with AP-1 and retinal developmental complexes

**DOI:** 10.1101/2025.09.04.674188

**Authors:** Edith De Bruycker, Olivier Zwaenepoel, George D. Moschonas, Delphine De Sutter, Eva D’haene, Charlotte Matton, Laure Vandevelde, Julie Van De Velde, Quinten Mahieu, Sara Dufour, Delphi Van Haver, Louis Delhaye, Simon Devos, Frauke Coppieters, Sven Eyckerman, Jan Gettemans, Elfride De Baere

## Abstract

The photoreceptor-specific nuclear receptor NR2E3 is a key transcription factor in the retinal transcriptional network, essential in photoreceptor cell fate and maintenance. Various biallelic loss-of-function variants lead to autosomal recessive inherited retinal disease (IRD), while a single missense variant G56R has been identified to cause autosomal dominant retinitis pigmentosa (adRP). However, a comprehensive understanding of the interactome of NR2E3 remains elusive. Here, we aimed to map the interactome of this nuclear receptor and the G56R pathogenic variant through proximity labelling followed by mass spectrometry. We used the TurboID “T2A split/link design” to identify the proximity interactome of wild-type NR2E3 and mutant G56R-NR2E3 in ARPE-19 cells. Several protein complexes involved in retinal development, including the AP-1 complex, were enriched. The results suggest similar interactomes for NR2E3 and G56R-NR2E3 in ARPE-19. However, BCOR was exclusively enriched in the G56R-NR2E3 network, strengthening the connection with the IRD phenotype. In conclusion, our approach has mapped the proximity interactome of NR2E3 and G56R-NR2E3, revealing interaction with protein complexes involved in (retinal) development.

## Introduction

Photoreceptor-specific nuclear receptor NR2E3, previously known as PNR, is a transcription factor in the retinal transcriptional network that plays a key role in photoreceptor cell fate and maintenance.^1^ It functions as a co-activator of rod-specific gene expression while concurrently suppressing cone-specific genes.^1, 2^ Biallelic loss-of-function variants in *NR2E3* (MIM #604485) have been associated with autosomal recessive inherited retinal diseases (IRD), including enhanced S-cone syndrome (ESCS)^3^, Goldmann-Favre syndrome (MIM #268100) and retinitis pigmentosa (arRP)^4^. A characteristic feature of recessive *NR2E3*-related disease is the excessive proliferation of cone photoreceptors derived from photoreceptor precursor cells (PPCs) that retain their default fate due to the loss of NR2E3, which normally suppresses S-cone gene expression during rod differentiation.^5^ In 2007, Coppieters *et al*. identified that NR2E3 variant p.(Gly56Arg) (p.(G56R), hereafter G56R), located in the DNA-binding domain, causes autosomal dominant retinitis pigmentosa (adRP).^6^ The glycine to arginine substitution in the first zinc finger motif ablates DNA binding, but does not hinder dimerization, thereby giving rise to a dominant-negative effect.^7, 8^ Moreover, *Nr2e3* has been shown to act as a modifier gene that rescues retinal degeneration and promotes homeostasis in RP models in mice.^9^

By means of yeast-two-hybrid (Y2H) and mammalian-two-hybrid (M2H) techniques several protein interaction partners of NR2E3 have been identified.^10–12^ In addition, through co-immunoprecipitation, the cone-rod homeobox (CRX) protein, neural retina leucine zipper (NRL) and NR1D1 were identified to co-regulate gene expression by NR2E3.^2^ Despite these studies, thus far only a limited number of interaction partners have been reported in comparison to other, more extensively studied nuclear receptors (NRs) such as the glucocorticoid (GR, *NR3C1*), androgen (AR, *NR3C4*) and estrogen receptors (ESR1, *NR3A1*). Therefore, we decided to further investigate and identify the complexes through which NR2E3 interacts to regulate gene expression. In addition, we were interested if a differential interactome could be observed for mutant G56R-NR2E3 compared to the wild-type NR2E3 protein.

In an effort to map the interactome of NR2E3 and G56R-NR2E3, we performed proximity labelling proteomics using the TurboID “T2A split/link design”^13^ in ARPE-19 cells. Our study revealed the presence of the HDAC3/NCoR1/2(SMRT) co-repressor complex known to interact with other/virtually all NRs. In addition, we identified the AP-1 complex that has been associated with retinal development and regulation of rod-specific genes in developing and adult rod photoreceptors. Moreover, we identified the PRC1 complex, a complex that is involved in transcriptional regulation of developmental genes. Mapping the proximal proteins of the G56R-NR2E3 counterpart revealed no outstanding differences in interactome composition compared to wild-type NR2E3, which is suggestive of an almost identical interactome in the cell type studied. As an exception, BCOR was significantly enriched in the G56R-NR2E3 network compared to wild-type NR2E3, strengthening the connection with the IRD phenotype.

## Results

### Generation of stable cell lines for proximity-dependent biotin labelling

Since there is no commercial human cell line available originating from rod-photoreceptor cells, the ARPE-19 cell line^14^ showing retinal precursor cell (RPC)-like behavior, was selected as a relevant model.^15^ For the biotin labelling we used TurboID, a hyperactive version of BioID.^16^ TurboID was genetically fused to the C-terminus of NR2E3 and G56R-NR2E3 with the ribosome skipping T2A peptide resulting in two independent proteins translated from a single mRNA transcript, and an inactivated version thereof as a linker sequence.^13, 17^ In the inactivated (MUTT2A) condition, ribosome skipping is abrogated and MUTT2A acts as a linker between the biotin ligase and NR2E3, resulting in the translation of a fusion protein. At the N-terminus of (G56R-)NR2E3 an HA-tag, and at the C-terminus of TurboID a V5-tag and nuclear localization signal (NLS) were introduced (Figure 1A). The NLS directs the biotin ligase towards the same subcellular localization as NR2E3 to improve the specificity of our control condition. (G56R-)NR2E3-T2A-TurboID and (G56R-)NR2E3-MUTT2A-TurboID were stably integrated in the genome of ARPE-19 cells under control of a constitutively active EF1α promotor using lentiviral transduction.

**Figure 1.**
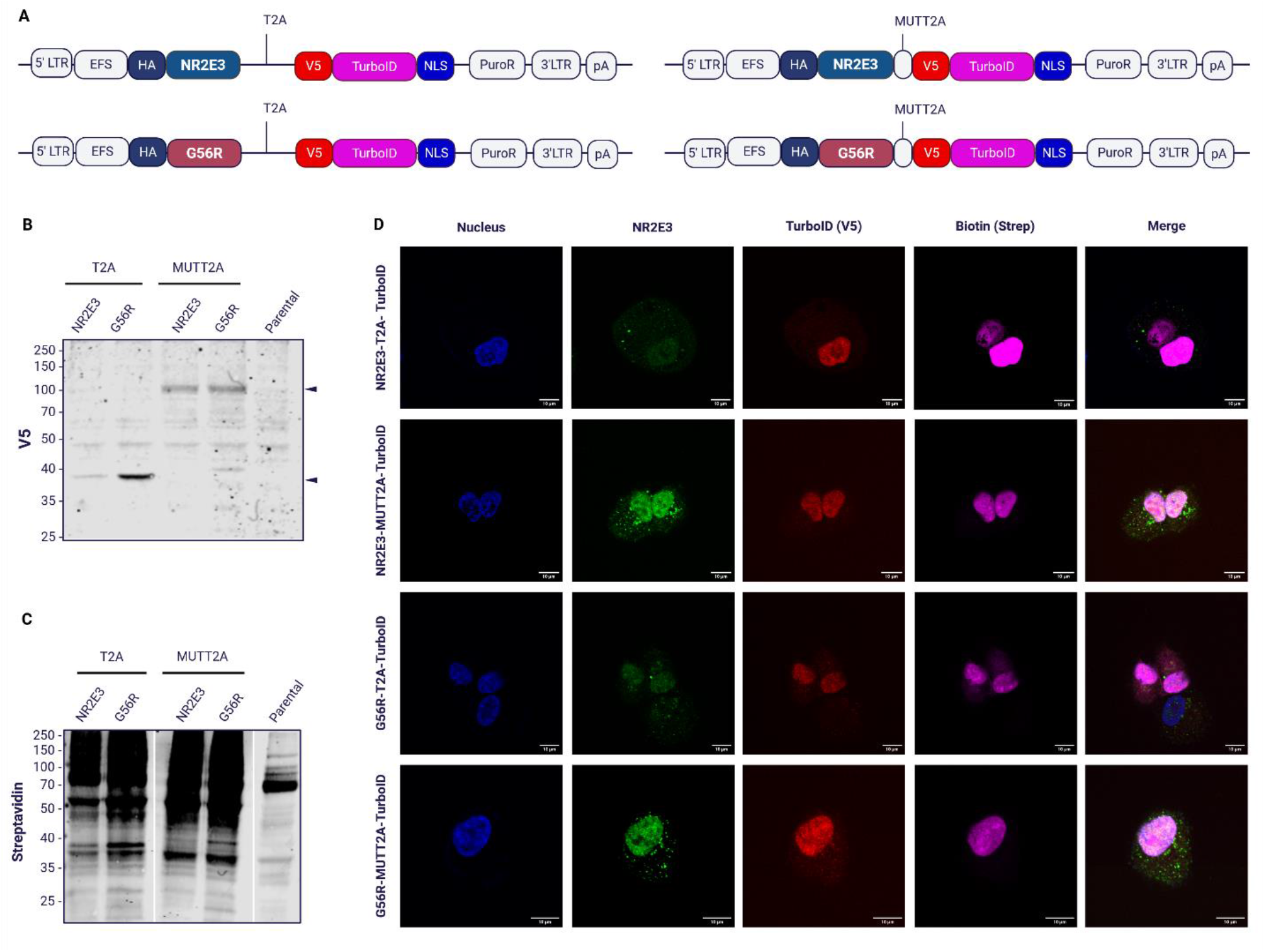
Generation of ARPE-19 stable cell lines for proximity-dependent labelling. **(A)** Schematic overview on the lentiviral constructs. NR2E3 (Blue) and G56R (Red) are genetically fused to TurboID at the C-terminus using the “T2A split/link design”. **(B)** Western blotting illustrates the expression of TurboID (38kDa) in the T2A condition and the fusion protein (85 kDa) in the MUTT2A condition. **(C)** Biotinylation (Streptavidin) staining of the NR2E3/G56R-(MUT)T2A-TurboID cell lines compared to non-transduced control. **(D)** Immunocytochemistry on transduced ARPE-19 cells expressing NR2E3/G56R (green) and V5-TurboID (red), and biotin staining (magenta).

Expression analysis of NR2E3 and G56R-NR2E3 through western blotting demonstrated the transcription of a single mRNA transcript translating into two distinct proteins in the T2A condition and into a fusion protein in the MUTT2A condition (Figure 1B). Upon treatment for 2 hours with biotin, a biotinylation pattern in all conditions could be observed (Figure 1C). Lastly, the expression, localization and biotinylation were evaluated through immunocytochemistry. In all four conditions, TurboID resides primarily in the nucleus showing a relevant subcellular localization (Figure 1D).

Proximity labelling was conducted in all four stable cell lines. Endogenously biotinylated proteins were enriched through streptavidin affinity purification and analyzed via LC-MS/MS with label-free quantification. A total of 2,407 proteins were recovered of which 2,261 proteins were reliably quantified. Principal component analysis (PCA) of all samples confirmed clustering of the T2A and MUTT2A conditions, but we did not observe distinct clustering of NR2E3 and G56R-NR2E3 samples (Supplementary Figure S1). This suggests that wild-type-NR2E3 and G56R-NR2E3 have a highly similar interactome in ARPE-19 cells.

### NR2E3 associates with multiple transcriptional regulatory complexes

A total of 51 proteins were significantly enriched for NR2E3. Several proteins involved in transcriptional processes and known to interact with NRs were identified in our dataset (Figure 2A), including nuclear receptor corepressor 1 (NCoR1) and NCoR2, also known as silencing mediator of retinoic acid and thyroid hormone receptor (SMRT) (Reactome ID: R-HSA-382093)^18^, and the class I histone deacetylase HDAC3. Together they form the co-repressor complex HDAC3/NCoR1/2, involved in primarily, but not exclusively, transcriptional repression (Figure 2B and C).^19, 20^ Nuclear receptor co-activator (NCoA), the molecular opposite of NCoR^21^, including NCoA1, was identified in our screen. While not significantly enriched for NR2E3, the coactivators have a log2FC greater than 0, suggesting a modest preference for NR2E3 compared to control. Members of the Polycomb repressive complex 1 (PRC1) PCGF1, USP34, RING1 and PCGF5 were significantly enriched for NR2E3.^22^ PRC1 complexes are composed of an E3 ubiquitin ligase (RING1A/B), the catalytic center of the complex, which interacts with one of six Polycomb group RING finger (PCGF) proteins. In our dataset, PCGF1 is highly enriched, which is known to form the PRC1.1 complex, a distinct subtype of PRC1, with RING1.^24^ Moreover, the PRC1.1 complex includes an additional ubiquitin ligase, SKP1. Though not statistically significant (p-val 0.08), SKP1 exhibits a log2FC above 2 (Figure 2A). This elevated fold change suggests a potential biological relevance.^22, 23^ Various subunits of the Activator Protein 1 (AP-1) transcription complex were significantly enriched. AP-1 is a collective term referring to homo-and heterodimers of the Jun, Fos, ATF and bZIP family.^24^ Our dataset revealed the presence of both the Jun and Fos family members. More specifically, JUN, JUNB and JUND were significantly enriched for the Jun family, and FOSL1 and FOSL2 for the Fos family. Furthermore, both Gene Ontology Enrichment Analysis (GOEA) and Spatial Analysis of Functional Enrichment (SAFE) reveal that a substantial subset of the enriched proteins are components of the spliceosome. In total, thirty out of 51 proteins were found to be correlated with the spliceosome, in particular, proteins from the mRNA cleavage and polyadenylation specificity factor (CPSF) complex constituted of five main subunits CPSF1, CPSF2, CPSF3, CPSF4 and FIP1L^25^, and the U2 small nuclear ribonucleoprotein (RNP) complex including SF3B1, SF3B3 and SF3B6 and SNRPA1 (Supplementary Figure S2). Lastly, several putative transcription factor binding sites (TFBSs) were enriched in the gene ontology (GO) analysis (TRANSFAC database) corresponding to EP300, NRF1 and PAX6 (Figure 2B).

**Figure 2.**
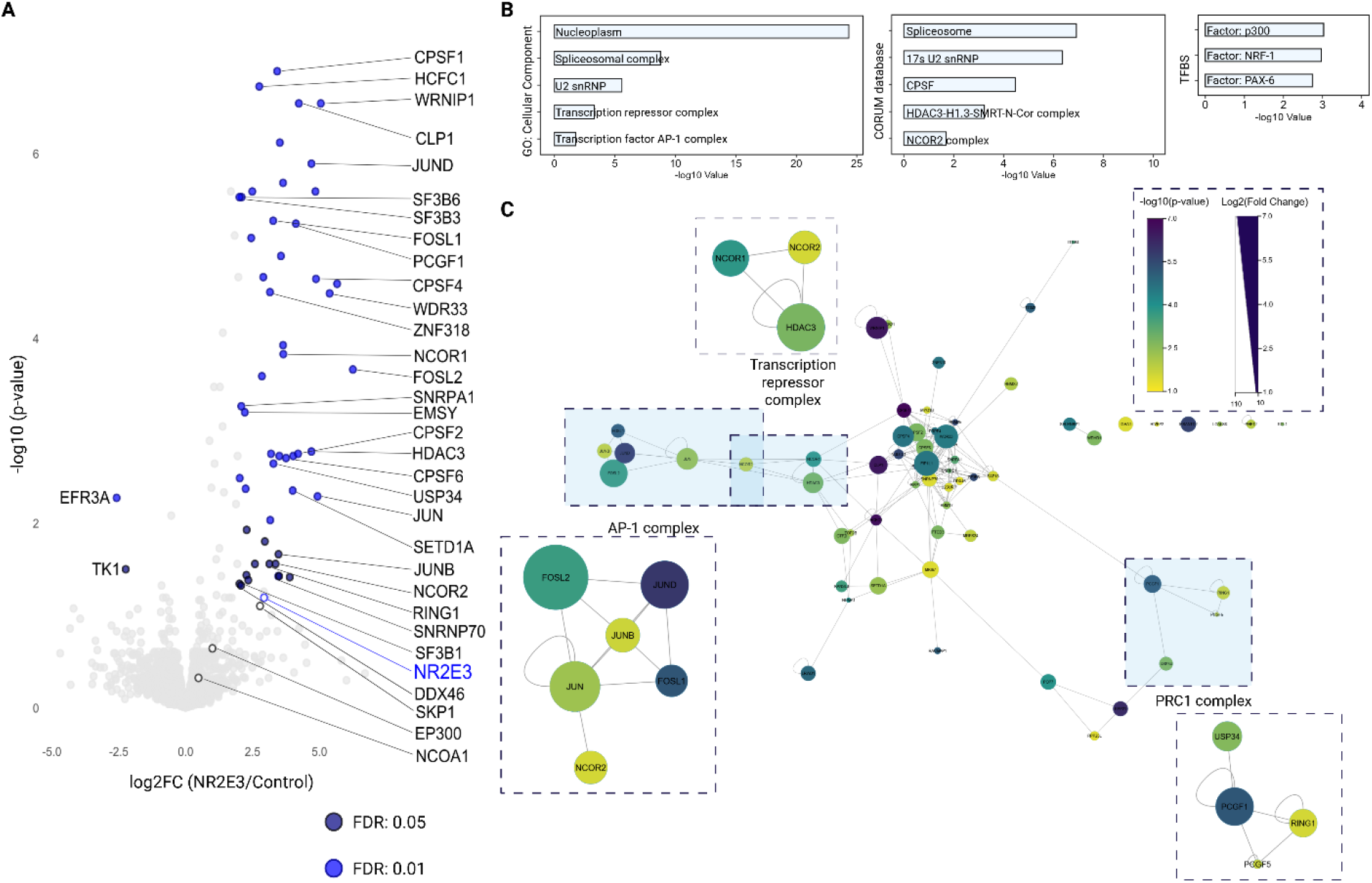
NR2E3 associates with transcriptional repressors in ARPE-19. **(A)** TurboID screen of NR2E3 in ARPE-19. A total of 51 potential interaction partners were enriched in the NR2E3-MUTT2A-TurboID dataset compared to T2A control. **(B)** Functional profiling of potential interaction partners of NR2E3. Significant proteins including proteins with moderate fold changes (log2FC>1, adjusted p-value<0.05) were taken into account to perform functional profiling in g:Profiler. Depicted are the results of GOEA for Cellular Component (CC), the CORUM database and putative TFBSs. **(C)** SAFE network visualizing core complexes found in our dataset corresponding to relevant GO enriched terms.^26, 27^ Each protein in the complex is represented with its corresponding p-value (-log10) and fold change (log2).

**Figure 3.**
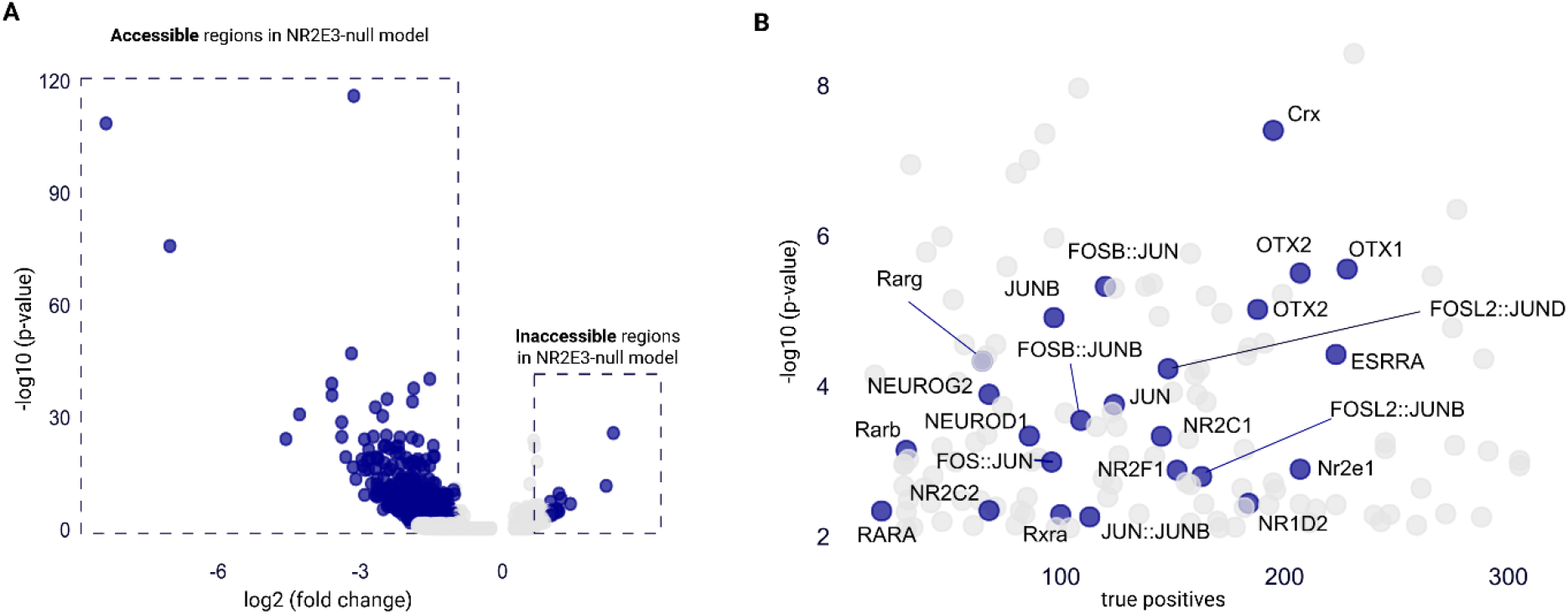
NR2E3 is primarily a transcriptional repressor. **(A)** Differentially accessible regions in *NR2E3*-null rod photoreceptors and isogenic controls. scATAC-seq data on retinal organoids^28^ reveals that 456 regions are accessible (log2FC<-1, p-value<0.05) in the *NR2E3*-null model and 13 peaks are inaccessible (log2FC>1, p-value<0.05) compared to the isogenic control. **(B)** Significantly accessible regions in the *NR2E3*-null model reveal transcription factor binding motifs. ATAC peaks that were accessible in rod photoreceptors of *NR2E3*-null retinal organoids reveal 131 known transcription factor biding motifs.

### scATAC seq data supports the association of NR2E3 with AP-1

Since loss of NR2E3 disrupts rod chromatin accessibility^28^, and our proteomic screen revealed its association primarily with transcription repressor complexes, we investigated if the single cell (sc)ATAC-seq data on retinal organoids^28^ supports the proximal interactome identified here. Regions with significantly altered chromatin accessibility of *NR2E3*-null rod photoreceptors compared to the isogenic controls were defined as those with a log2FC greater than 1 (inaccessible in *NR2E3*-null) or less than −1 (accessible in *NR2E3*-null), with a p-value lower than 0.05. This identified 456 significant accessible peaks and 13 significant inaccessible peaks in the *NR2E3*-null model (Figure 2D). Within the ATAC peaks that were accessible in the *NR2E3*-null rod-photoreceptors, simple enrichment analysis (SEA)^29^ was performed, identifying 131 significant transcription factor binding motifs (Figure 2E). Among the strongest hits, we find Orthodenticle Homeobox 1 and 2 (OTX1/2), and cone-rod homeobox protein (CRX), which are transcription factors essential to retina neurogenesis and photoreceptor differentiation.^30–34^ In addition, the analysis reveals transcription factor binding motifs of several components of the AP-1 complex including JUN, JUNB, JUND and FOSL2, further supporting an association of these proteins with NR2E3, as evidenced by our BioID screen.

### AP-1 is specific to NR2E3

The interactomes of other, more extensively studied NRs (Figure 4A) and NRs involved in retinal development (Figure 4B), were compared to the NR2E3 dataset. The enriched proteins for NR2E3 were compared to the interactome, as indicated in BioGRID, of the three extensively studied steroid receptors, showing a large overlap with the interactome of NR3C4 (53%) and NR3A1 (47%) (Figure 4A). Four proteins show a shared interaction between all four NRs. The two NRs playing a role in the retina show an overlap of solely three and six proteins with NR2E3, respectively, corresponding to the HDAC3/NCoR1/2 co-repressor complex (Figure 4B). Furthermore, we extended our analysis by comparing the significantly enriched proteins with the interactome of all 47 other NRs and the Contaminant Repository for Affinity Purification (CRAPome) (Figure 4C). The CRAPome is a repository that compiles negative controls from BioID studies, enhancing the identification of true interaction partners while minimizing background noise.^35^ A high frequency of detection (FoD) value indicates that the protein is frequently detected across experiments. The proteins involved in splicing are frequently enriched in BioID experiments according to the CRAPome database, with an FoD of approximately 50% for the members of the SF3b complex (SF3B1, SF3B3 and SF3B6 and SNRPA1). This suggests that these results should be interpreted with caution. Furthermore, the analysis illustrates that the HDAC3/NCoR1/2 complex has been experimentally validated as being associated with approximately 50% of all NRs. In contrast, the analysis demonstrates that most of the AP-1 complex components are highly specific NR2E3 interactors.

**Figure 4.**
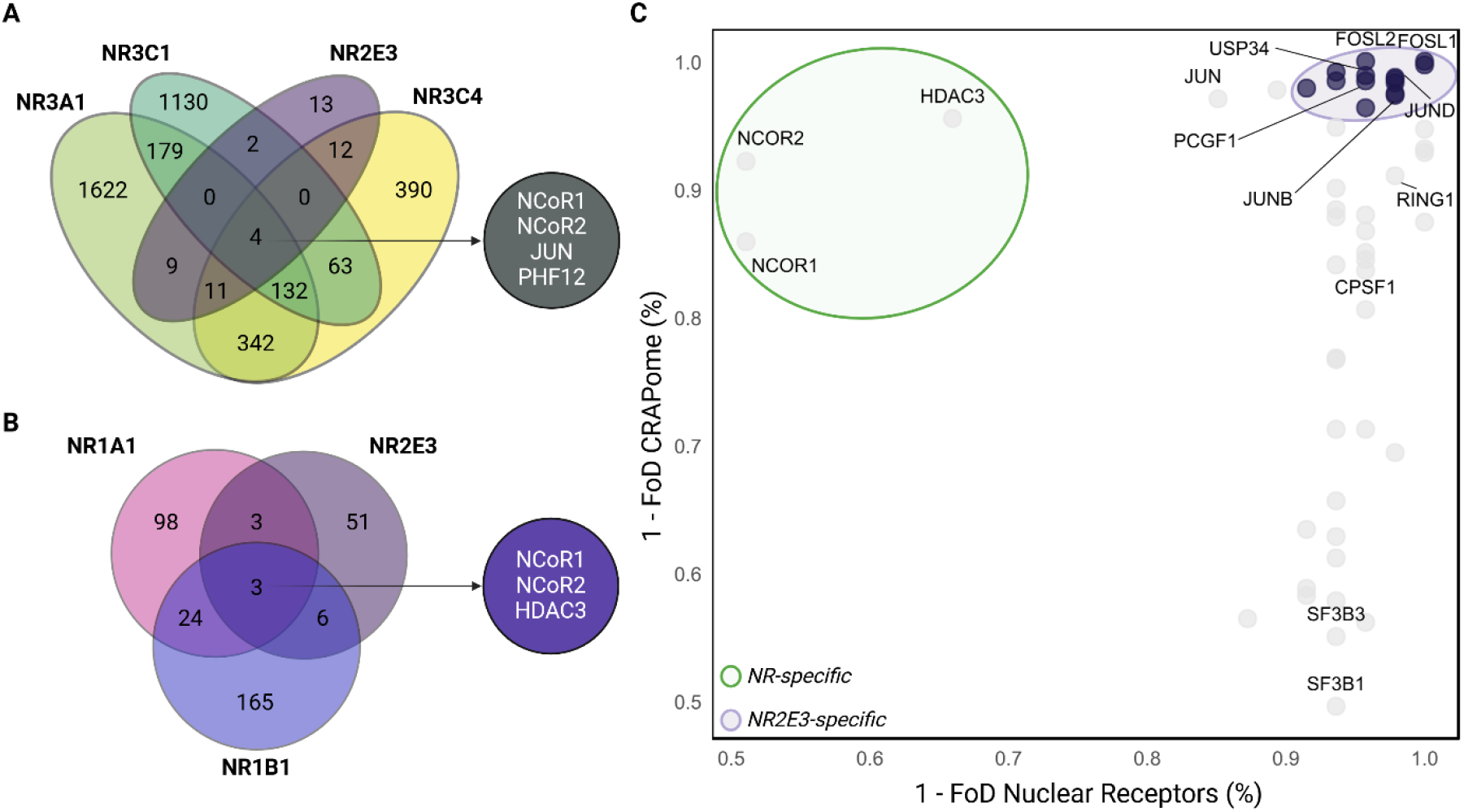
The AP-1 complex is highly specific to NR2E3. (**A**) Comparison of enriched partners for NR2E3 with extensively studied NRs NR3A1 (ESR1), NR3C4 (AR) and NR3C1 (GR). **(B)** Venn diagram displaying the number of interactors overlapping with NRs involved in retinal development: NR1B1 (RARA) and NR1A1 (THRA). **(C)** Frequency of Detection (FoD) in the interactome of all NRs and the CRAPome. Proteins highlighted in blue are detected in fewer than 5% and 10% of BioID experiments (CRAPome) and NR interactomes, respectively.

### Bcl-6 co-repressor (BCOR) is exclusively enriched for G56R-NR2E3 in an otherwise highly similar interactome to wild-type NR2E3

Differential analysis using stringent cut-off values (FDR 0.05) showed 61 proteins to be significantly enriched for G56R-NR2E3 compared to the MUTT2A control (Figure 5A). Forty-four of these interactors overlapped with NR2E3 (Figure 5B), while the majority of the remaining 17 proteins exhibited only minor differences in fold change and/or adjusted p-value, despite not reaching significance in the NR2E3 dataset (Figure 5C). The three co-repressor complexes HDAC3/NCoR1/2, PRC1.1 and AP-1 were similarly significantly enriched, suggesting a highly comparable interactome to NR2E3. However, Bcl-6 co-repressor (BCOR), a member of the PRC1.1 complex, is exclusively enriched for G56R-NR2E3 (FDR=0.01) (Figure 4E).

**Figure 5.**
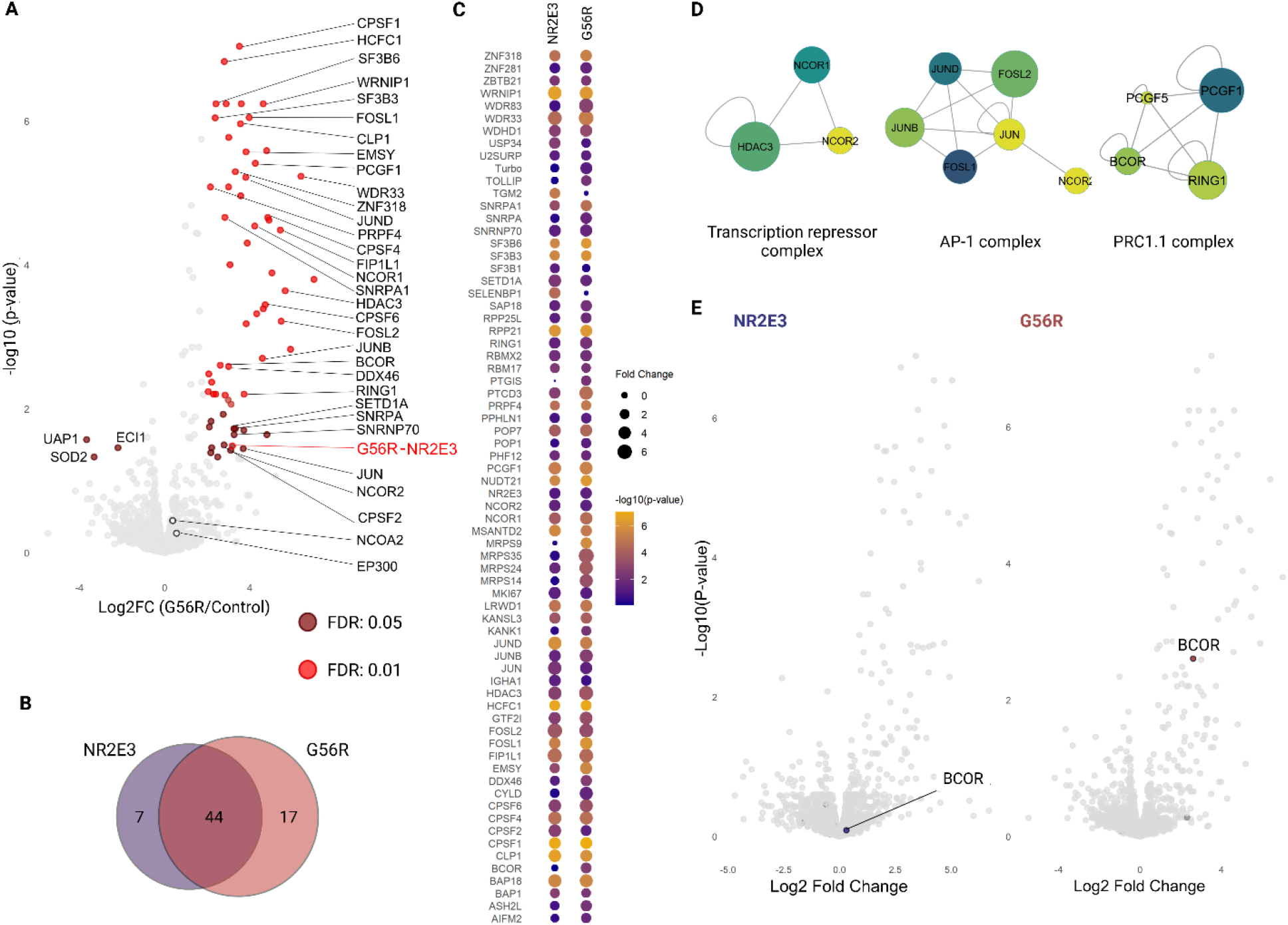
TurboID on G56R-NR2E3 demonstrates a highly similar interactome compared to NR2E3 in ARPE-19. (**A**) Enriched proteins for the G56R mutant. A total of 61 proteins were significantly enriched proteins for G56R compared to T2A control. (**B**) The proximal interactomes of G56R-NR2E3 and NR2E3 show a high overlap. A total of 44 significantly enriched proteins can be detected in both datasets. (**C**) All significantly enriched proteins for both datasets. A total of 68 proteins are significantly enriched in at least one dataset. Differences can be observed for adjusted p-values (color) and log2 fold changes (size). (**D**) The biologically relevant complexes for NR2E3 are enriched for G56R-NR2E3. (**E**) BCOR is exclusively enriched for G56R-NR2E3.

## Discussion

In this study, the proximal interactome of wild-type photoreceptor-specific nuclear receptor NR2E3 and mutant G56R-NR2E3 were mapped in ARPE-19 cells using the TurboID “T2A-split/link design”.^13^ TurboID is a robust technique for investigating protein-protein interactions (PPIs) within living cells.^16, 36^ This method leverages proximity-dependent protein labelling, wherein a promiscuous biotin ligase is fused to the protein of interest.^37^ Upon exogenous addition of biotin to the cell culture medium, the fusion protein facilitates the biotinylation of proteins in close vicinity. This method enables the identification of direct, indirect and transient interaction partners.^36^ The use of T2A^17^ allows for the translation of two distinct proteins from a single mRNA transcript, thereby resulting in the simultaneous expression of (G56R-)NR2E3 and TurboID in the control set-up. This control cell line undergoes the same changes introduced by the presence of (G56R-)NR2E3, preventing control bias, since the introduction of a protein may impact the global proteome, especially in the case of transcription factors. By mutating the T2A sequence (MUTT2A), ribosome skipping no longer occurs, resulting in the translation of the fusion protein.^38^ This strategy has proven effective for other NRs enhancing the understanding of their functions.^39^

A total of 51 and 61 proteins were significantly enriched for NR2E3 and G56R-NR2E3, respectively. Three transcriptional co-repressors were identified in both screens, including the HDAC3/NCoR1/2 (SMRT), PRC1.1 and AP-1 complex. Given that HDAC3/NCoR1/2 is a well-established interactor of NRs, this complex represents a robust positive control within our BioID screen. Moreover, this complex has been shown to interact with NR2E3 in a ligand-independent state, thereby mediating transcriptional repression.^8, 40^ Histone deacetylase 3 (HDAC3) is a member of the HDAC Class I family of epigenetic regulators and exerts its deacetylase activity by forming a multiprotein complex with Nuclear Co-repressor 1 and 2 (NCoR1/2).^41^ While the role of the HDAC3/NCoR1/2 complex in the retina remains largely unexplored, it is known to regulate genes involved in development, metabolism and circadian rhythm in various tissues, including neuronal tissue.^18, 19, 42, 43^ Furthermore, the circadian rhythm is largely regulated by HDAC3 in complex with Rev-Erbα (NR1D1), which has been shown to interact with NR2E3 through recombinant pull-down assays.^2, 44^ However, we did not detect NR1D1 in our dataset. While the protein is expressed in ARPE-19 cells^45^, previous shotgun proteomics on ARPE-19^46^ (PXD022545) also failed to identify it. This suggests that NR1D1 may not be easily detectable through LC-MS/MS and be a potential false negative in our data.

Members of the Polycomb group Repressive Complex 1 (PRC1) complex but not PRC2 were enriched in the NR2E3 dataset. Importantly, PRC1 is involved in chromatin compaction resulting in gene repression in a PRC2-independent manner.^22, 47, 48^ In addition, this complex plays a key role in regulating primarily homeobox genes during development through ubiquitination of histone 2A (H2A), which is a marker for gene silencing.^49^ The recruitment of PRC1 to specific chromatin sites is influenced by the composition of accessory proteins.^48^ This suggests that NR2E3 may act as an accessory protein in a rod-specific context, facilitating chromatin compaction and, in turn, silencing of cone-specific genes. Surprisingly, BCL6 co-repressor (BCOR), a Polycomb repressive complex 1 factor, was significantly enriched in the G56R-NR2E3 dataset but not in NR2E3.^23, 50^ BCOR is a co-repressor of CRX and OTX2 involved in retinal development.^51^ Furthermore, the co-repressive activity of BCOR on CRX/OTX2 is essential for regulating photoreceptor gene expression levels, thereby safeguarding photoreceptor survival. Interestingly, missense variants in BCOR have been linked to severe early-onset X-linked retinal degeneration.^51^ Therefore, if BCOR interacts with G56R-NR2E3, this interaction could impair the ability of BCOR from binding DNA, potentially intensifying retinal degeneration in NR2E3-associated adRP.^51^ Lastly, an important interactor of the PRC1 complex in photoreceptors is Sterile alpha motif domain containing 7 (SAMD7) of which an interaction with NR2E3 has been established in HEK293 cells upon overexpression.^49, 52^ In addition, SAMD7 functions as a CRX-regulated transcriptional repressor in the retina, with bi-allelic pathogenic variants linked to IRD.^53, 54^ However, due to its neural retina-specific expression, we could not verify this interaction in ARPE-19. Both shotgun proteomics (PXD022545) and RNA seq (GSE88848) of ARPE-19 could not detect expression of SAMD7 in ARPE-19. This suggest that SAMD7 is not expressed in ARPE-19 and cannot be detected in our screen.

The third co-repressor complex enriched in our data is activator protein-1, a collective term referring to transcription factor dimers composed of members from the Jun and Fos family. AP-1 is an essential complex involved in virtually all cellular processes through remodeling chromatin at distal enhancers harboring the AP-1-binding site.^24, 55^ It has been described that AP-1 is involved in retinal development through the interaction with NRL.^56, 57^ In addition, the promotor region of rod-specific genes and opsins harbor AP-1 response elements, implying that the AP-1 complex is involved in the regulation of photoreceptor genes.^58^ Other NRs have also been shown to interact with AP-1^59, 60^, by which AP-1 mediates recruitment of the NR towards regulatory elements and facilitate interactions with other co-factors. Importantly, the majority of the Jun and Fos family members (JUNB, JUND, FOSL1, FOLS2) are almost exclusively represented within the NR2E3 dataset, in comparison to the interactome of all other NRs. Furthermore, this interaction could be confirmed through scATAC-seq data comparing chromatin accessibility of *NR2E3*-null retinal organoids with a wild-type isogenic control published by Mullin, *et al*. (2024).^28^ The predominance of the gained accessibility in the *NR2E3*-null model indicates that NR2E3 functions primarily as a global repressor.^28^ Previous studies have demonstrated that CRX interacts with NR2E3 and co-occupies promotor and enhancer regions, thereby activating and repressing rod-specific and cone-specific genes, respectively.^2, 61^ Therefore, the scATAC-seq results may be suggestive of potential transcription factor co-occupancy. Motif enrichment analysis of the repressed regions by NR2E3 revealed the presence of AP-1 binding sites, which could also confirm the co-occurrence of NR2E3 with the AP-1 complex in mediating gene repression of developmental genes and further supporting the interaction of this complex with NR2E3.

Furthermore, TRANSFAC revealed the presence of EP300, NRF1 and PAX6 in the enriched dataset for NR2E3. This entails that the enriched proteins for NR2E3 either bind the transcription factor binding sites or are downstream targets of these genes. Histone acetyltransferase p300 is an important co-activator of several NRs including the glucocorticoid, estrogen and androgen receptor.^62–66^ It was suggested that p300 may be involved in photoreceptor maturation and maintenance as well as expression of key rod-specific genes. Nuclear respiratory factor 1, NRF1, is a transcriptional regulator (activator) of nuclear genes essential to mitochondrial biogenesis and respiratory factors with a major role in neurons.^67–69^ It has been demonstrated that NRF1 is expressed in proliferating retinal progenitor cells (RPCs) and enriched in cells with high energy demands, including rods.^70^ *Nrf1* deficiency affects retinal development causing thinner retinas, eventually resulting in retinal degeneration.^70^ Lastly, Paired box protein (PAX) is a transcription factor family involved in early development^71^, with PAX6, also known as oculorhombin, primarily involved in the development of the central nervous system including the retina.^72, 73^ Moreover, PAX6 maintains the multipotency of RPCs and orchestrates photoreceptor differentiation at the appropriate time point.^74^

Our dataset revealed the presence of two large complexes involved in RNA maturation. The complete CPSF complex constituted of CPSF1-6, WDR33^75^ and FIP1L^25^, involved in 3’ end capping of pre-mRNA, was significantly enriched.^25, 76^ In addition, the SF3b complex (SF3B1/3/6 and SNRPA1) which is part of the U2 snRNP complex, was enriched. This complex is involved in pre-spliceosomal assembly through binding with the branch point sequence and sequestering the U4/U6-U5 tri snRNP complex.^77–79^ Furthermore, a physical interaction between these two complexes through SF3B3 and CPSF1 has been demonstrated, which is essential to splicing.^80^ Intriguingly, pathogenic variants in ubiquitously expressed splicing factors have been associated repetitively with retina-specific diseases, including adRP.^81, 82^ Therefore, if NR2E3 plays a role in the spliceosome, mutations in this gene potentially contribute to retinal degeneration due to the sensitivity of the retina to alterations in the spliceosome. Importantly, the CRAPome (Figure 4C) illustrated that the spliceosome is frequently enriched (approx. 50% of BioID experiments). Therefore, further validation is required to demonstrate the potential link with the spliceosome.

In conclusion, absence of significantly enriched co-activator complexes in our data suggests that NR2E3 is primarily a transcriptional repressor. The data presented by Mullin *et al*. (2024)^28^ supports the repressor function. In addition, NR2E3 has been shown to mediate transcriptional repression of developmental genes through its interaction with the AP-1 complex. Furthermore, our data suggested an interaction with the PRC1 complex, of which BCOR was exclusively enriched in the G56R-NR2E3 dataset. This might serve as another piece of the puzzle contributing to the phenotype associated with adRP.

## Materials & Methods

### Stable cell line generation for proximity-dependent labelling and affinity purification

Four ARPE-19 cell lines expressing either wild-type or mutant NR2E3-(MUT)T2A-TurboID were generated by transduction with lentivirus. The lentiviral constructs were manufactured according to an established protocol.^13^ In brief, an *in-house* golden gate modular cloning platform was employed to assemble different subunits (Level-0 constructs) simultaneously into a single transcriptional unit (Level-1 construct), followed by the assembly of two transcriptional units into a single lentiviral entry vector (Level-2 construct) expressing the transgene and a selectable marker. Four plasmids were generated expressing NR2E3-(MUT)T2A-TurboID or G56R-(MUT)T2A-TurboID and puromycin resistance gene. MUTT2A has the phenylalanine to proline substitution in the T2A sequence at position 18 (T2A_p.F18P). Additionally, NR2E3(-G56R) and TurboID were N-terminally tagged with HA and V5, respectively. At the C-terminus of TurboID, an additional SV40 nuclear localization signal (NLS) (SRADPKKKRKV) was introduced to correct for the subcellular localization of the control set-up.

For the lentivirus production, 2.2×10^6^ HEK293T cells were seeded for each set-up. At 75-100% confluency, cells were co-transfected by Calcium Phosphate Precipitation with the Level-2 construct, pCMV-deltaR8.2 (Addgene#12263) and pMD2.G (Addgene#12259). Twenty-four hours post transfection, cells were washed once with DPBS (Catalog No. 12037539, Fisher Scientific Belgium) and fresh culture medium supplemented with 10% FBS was administered onto the cells. Lentivirus-containing medium was collected at 24 and 48 hours post-transfection, after which cellular debris was removed by centrifugation for 3 min at 2,000 rpm, followed by filtration through a 0.22 µm filter. Crude supernatant was stored at −80°C until transduction or used immediately after collection.

For each stable cell line, ARPE-19 cells were seeded in a 6-well plate and transduced with the respective lentivirus at low MOI. Polybrene was added to all transductions at a final concentration of 8 µg ml^−1^. Twenty-four hours post transduction, medium was refreshed. Transduced ARPE-19 cells were selected through antibiotic selection with puromycin at a final concentration of 1 µg ml^−1^ (Catalog code ant-pr-1, Invivogen). After at least one week of selection and until non-transduced control cells were completely eliminated under selection conditions, cells were expanded for downstream experiments.

The four stable cell lines were cultured in growth medium composed of Dulbecco’s Modified Eagle Medium: Nutrient Mixture F12 (DMEM-F12, Catalog No. 11540446, Life Technologies Europe), supplemented with GlutaMAX (Catalog no. 12077549, Fisher Scientific Belgium), 10% fetal bovine serum (Catalog no. A5256701, ThermoFisher Scientific), 1 µg ml^−1^ puromycin, 100 units ml^−1^ penicillin and 100 units ml^−1^ streptomycin (Catalog No. 11548876, Fisher Scientific Belgium). Cells were maintained at 37°C in 5% CO_2_.

### Immunofluorescence staining

For each set-up, 5×10^4^ cells were seeded on a cover slip and incubated for at least 24 hours until immunofluorescence staining. Cells were fixed in 3% paraformaldehyde, permeabilized with 0.1% Triton X-100 and neutralized in 50 mM glycine. In 1% of bovine serum albumin (Catalog No. A2153, Sigma), samples were blocked and incubated with primary antibody anti-V5 (Catalog No. 10347642, Invitrogen) and anti-NR2E3 (Catalog No. 14246-1-AP, Proteintech Europe) for 1h at 37°C and Alexa-conjugated secondary antibody for 30 min at room temperature. Nuclei were stained with DAPI at a final concentration of 0.4 µg ml^−1^ and the target proteins using Alexa-Fluor488 (Catalog No. 10236882, Invitrogen) and Alexa-Fluor-594 (Catalog No. 10644773, Invitrogen). Cells were mounted using VectaShield Mounting Medium (Catalog No. H-1000-10, Vector Laboratories). For imaging, the confocal laser scanning microscope (FV1000, Olympus) was used.

### BioID

Proximity-dependent biotin labelling was performed on all stable ARPE-19 cell lines simultaneously. For every cell line, four replicates were taken along, resulting in four T2A and four MUTT2A samples for both NR2E3 and G56R-NR2E3. For each replicate, 5×10^6^ cells were seeded on a 150 mm culture dish. Forty-eight hours after seeding, proximity labelling by TurboID was initiated by adding 50 µM biotin (Catalog No. B4639, Sigma).^16, 83^ After two hours of biotinylation, cells were washed twice in DPBS and lysed in a total volume of 750 µl RIPA buffer (150 mM Sodium Chloride, 50 mM Tris-HCl at pH 8, 1% NP-40, 0.5% Sodium Deoxycholate, 0.5% Sodium Dodecyl Sulphate (SDS), 1mM EDTA) supplemented with 1 mM PMSF (Phenylmethylsulfonyl fluoride) and 200 µg ml^−1^ Protease-inhibitor cocktail. Lysates were incubated for 1 hour at 4°C over agitation with 250 U Benzonase® Nuclease (Catalog No. E1014, Merck) to reduce the viscosity of the samples. To remove debris, lysates were centrifuged for 15 minutes at 14,000 x *g* at 4°C and supernatant was transferred to a new tube. Total protein concentration was measured with the Protein Assay (Catalog No. #5000006, Bio-Rad) across all samples and the maximum shared protein amount was used as input material for the pull-down. For every sample, 30 µl of Streptavidin Sepharose™ High Performance beads (Catalog No. GE17-5113-01, Cytiva) was prepared for biotin enrichment by three washing steps in RIPA buffer and centrifugation at 400 x *g* for 1 minute. Affinity purification was performed over agitation at 4°C for 3 hours after which beads were recovered through centrifugation at 400 x *g* for 1 minute at 4°C. Three stringent washing steps in RIPA buffer were performed and twice in 50 mM bicarbonate (pH 8). Proteins were eluted in 50 mM Triethylammonium bicarbonate buffer (TEAB) with 5% SDS.

### Sample handling

Tris-(2-Carboxyethyl)phosphine (TCEP) and Chloroacetamide (CAA) were added to final concentrations of 10 mM and 40 mM, respectively, to reduce disulfide bridges in cysteine residues and to alkylate cysteine residues. Samples were incubated for 10 minutes in the dark at 95°C followed by acidification by adding phosphoric acid to a final concentration of 2.75%. Next, samples were neutralized in wash buffer (100 mM TEAB (pH 7.55), 90% MeOH), transferred to a Micro S-trap and concentrated through centrifugation at 4,000 *x g*, followed by three wash steps in wash buffer. Receiver tube was replaced by a protein LoBind Eppendorf. Overnight trypsinization was performed at 37°C in 50 mM TEAB buffer with 1 µg of trypsin. For mass spectrometry analysis, in 50 mM TEAB buffer 50% acetonitrile (ACN) was added, followed by centrifugation for 1 minute at 4,000 x *g*, and supplemented with 0.2% FA. Peptide mixtures were dried by speedvac evaporation until analysis.

### Liquid chromatography-mass spectrometry

Peptides were re-dissolved in 20 µl loading solvent A (0.1% trifluoroacetic acid in water/ACN (99.5:0.5, v/v)) of which 2 µl of sample was injected for LC-MS/MS analysis on an ultimate 3000 ProFlow nanoLC system in-line connected to a Q-Exactive HF mass spectrometer (Thermo) equipped with pneu-Nimbus dual ion source (Phoenix S&T). Trapping was performed at 20 μl minute^−1^ for 2 minutes in loading solvent A on a PepMap™ Neo Trap column (Thermo Scientific, 300 μm internal diameter (I.D.), 5 μm beads). The peptides were separated on 250 mm Odyssey Ultimate, 1.7 µm C18, 75 µm inner diameter (Ionopticks) kept at a constant temperature of 45°C.

Peptides were eluted by a gradient reaching 26.4% MS solvent B (0.1% FA in ACN) after 30 min, 44% MS solvent B at 38 min, 56% MS solvent B at 40 min followed by a 5 minute wash step at 56% MS solvent B and re-equilibration with MS solvent A (0.1% FA in water). The flow rate was set to 250 nl minute^−1^.

The mass spectrometer was operated in data-independent (DIA) mode, automatically switching between MS and MS/MS acquisition. Full-scan MS spectra ranging from 375-1500 m/z with a target value of 5E6, a maximum fill time of 50 milliseconds and a resolution of 60,000 were followed by 30 quadrupole isolations with a precursor isolation width of 10 m/z for HCD fragmentation at an NCE of 30% after filling the trap at a target value of 3E6 for maximum injection time of 45 milliseconds. MS2 spectra were acquired at a resolution of 15,000 in the Orbitrap analyser. The isolation intervals ranging from 400 – 900 m/z, without overlap were created with the Skyline software tool. The polydimethylcyclosiloxane background ion at 445.120028 Da was used for internal calibration (lock mass) and QCloud has been used to control instrument longitudinal performance during the project.^84, 85^

### Data analysis

Analysis of the MS data was performed in DiaNN (version 1.9.2)^86^. Precursor false discovery rate (FDR) was set at 1%. Spectra were searched against human protein sequences in the UniProt database (9606). Protein sequences of HA-NR2E3-T2A, HA-G56R-T2A and V5-Turbo-NLS were added to the search. Enzyme specificity was set as C-terminal to arginine and lysine, also allowing cleavage at proline bonds with a maximum of 1 missed cleavage. Fixed modification was set to carbamidomethylation of cysteine residues. Matching between runs was enabled. Mainly default settings were used, except for the addition of a 400-900 m/z precursor mass range filter and MS1 and MS2 mass tolerance was set to 10 and 20 ppm, respectively. Further data analysis of the results was performed with an in-house script in the R programming language. Protein expression matrices were prepared as follows: the DIA-NN main report output table was filtered at a precursor, and protein library q-value cut-off of 1%, and only proteins identified by at least one proteotypic peptide were retained. After pivoting into a wide format, iBAQ intensity columns were added to the matrix using the DIAgui’s R package get_IBAQ function.^87^ PG MaxLFQ intensities were log2 transformed and replicate samples were grouped. Proteins with less than 3 valid values in at least one group were removed and missing values were imputed from a normal distribution centred around the detection limit (package DEP)^88^, leading to a list of 2,261 quantified proteins in the experiment. To compare protein abundance between pairs of sample groups NR2E3-G56R-mT2A-TurboID vs NR2E3-G56R-T2A-TurboID and NR2E3-mT2A-TurboID vs NR2E3-T2A-TurboID, statistical testing for differences between two group means was performed, using the package limma^89^. Statistical significance for differential regulation was set to a false discovery rate (FDR) of < 0.05 and |log2FC| = 2. Z-scored PG MaxLFQ intensities from significantly regulated proteins were plotted in a heatmap after non-supervised hierarchical clustering.

### Motif enrichment analysis from scATAC-seq data

Single-cell (sc)ATAC-seq peak data from NR2E3-null retinal organoids and isogenic controls were retrieved from Mullin *et al*. (2024).^28^ Differentially accessible regions (DARs) were filtered for a |log2FC| higher than 1 and p-value (p-val) less than 0.05. Sequences corresponding to these DARs were extracted and formatted to FASTA format using BEDTOFASTA in MEME Suite (version 5.5.7) (https://meme-suite.org), with reference to the hg38 human genome from UCSC Mammal Genomes. To identify transcription factor binding motifs enriched in the significantly accessible regions of the NR2E3-null model, Simple Enrichment Analysis (SEA)^29^ algorithm was applied with the following parameters: sea --verbosity 1 --oc. --thresh 10.0 --align center --p peak_seq.FASTA --m motif_db/JASPAR/JASPAR2022_CORE_vertebrates_redundant_v2.meme.

### Frequence of Detection (FoD)

The significantly enriched proteins were queried in the CRAPome database against the *H. sapiens* – Proximity Dependent Biotinylation Database encompassing 716 experiments. For each protein the frequency of detection (%) was extracted by dividing the number of times the protein was found in an experiment by the total amount of experiments in the database. Furthermore, the enriched proteins were compared to all known interaction partners of the NR, thereby creating a FoD for NRs specifically. The 1-FoDs from the CRAPome and NRs were projected on the y-axis and x-axis, respectively.

## Acknowledgements

E.D.Ba. acknowledges support by the Ghent University Special Research Fund (BOF20/GOA/023); EJPRD19-234658 Solve-RET (E.D.B.); Research Foundation Flanders FWO (1802220N). E.D. and C.M. acknowledge support by an FWO post-doctoral (12D8523N) and PhD fellowship (11Q3D24N) respectively. L.D. acknowledges support by a UGent BOF post-doctoral mandate (BOF22/PDO/024). SE thankful for a FWO project grant (G042918N). We would like to thank the VIB Proteomics Core for their guidance and operating the mass spectrometer. E.D.Ba. is member of ERN-EYE (Framework Partnership Agreement No 739534-ERN-EYE).

## Author contributions

E.D.Br. conceptualized the original idea of this study and the experimental design. G.D.M., S.E., L.D., D.V.H. and S.D.V. contributed to the experimental design. D.D.S. and S.E. provided the constructs for lentivirus production. E.D.Br. performed all wet-lab experiments. O.Z., J.V.D.V., Q.M. and L.V. assisted with laboratory work. S.D.V. supervised the mass-spectrometry analysis. S.D.F. and E.D.Br. performed the proteomics data analysis. G.D.M., and L.D., assisted with data analysis. E.D.H. and C.M. assisted with the re-analysis of public datasets. E.D.Ba., and J.G. supervised the project. E.D.Br. wrote the original draft of the manuscript. E.D.Ba., and J.G. and F.C. edited the original draft of the manuscript. All authors edited and contributed to the final manuscript.

## Disclosure and competing interests statement

The authors declare no competing interests.

## Data availability section

The mass spectrometry proteomics data have been deposited to the ProteomeXchange Consortium via the PRIDE^90^ partner repository with the dataset identifier PXD068057.

## Supplementary figures

**Figure S1.**
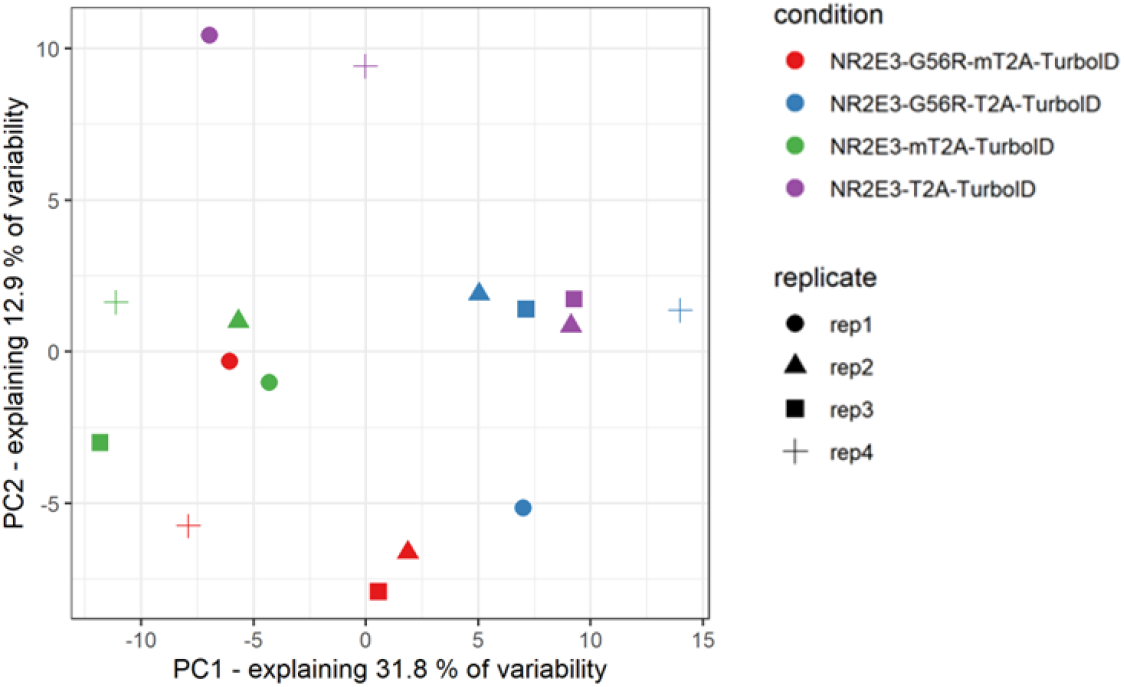
Principal component analysis of the BioID samples. The PCA plot illustrates sample mappings along the two principal components (PCs). Percentages of explained data variance for each PC are shown on the x and y axis.

**Figure S2.**
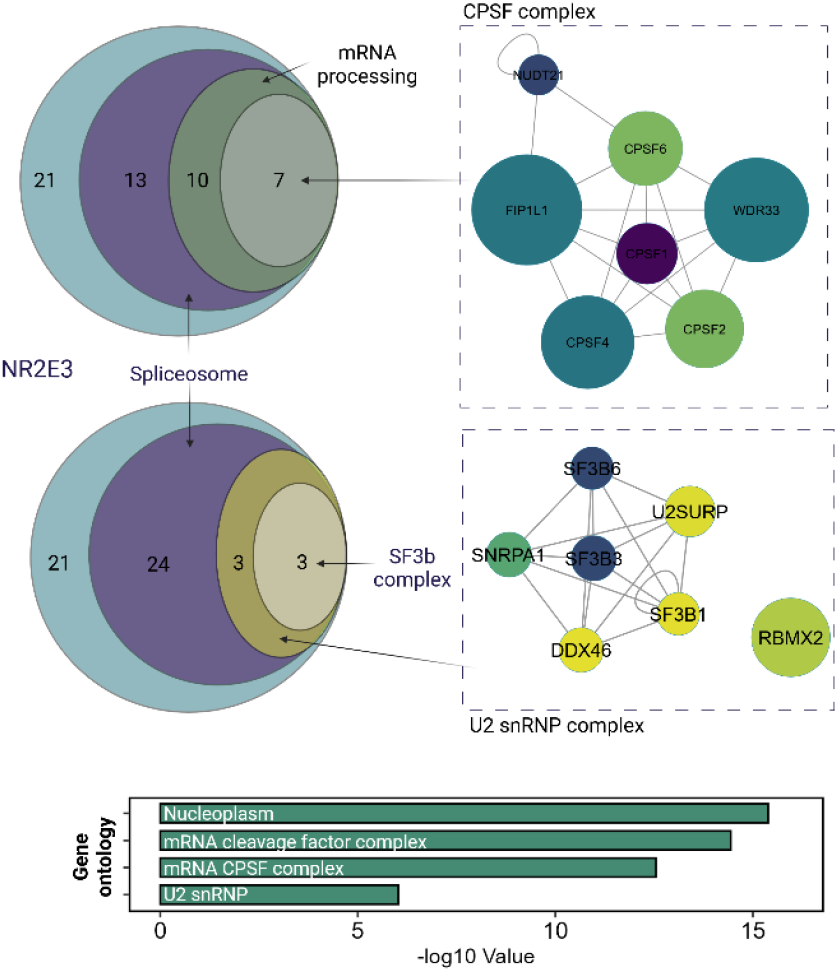
The enriched proteins for NR2E3 reveal a potential connection with the spliceosome. Thirty out of 51 of the enriched proteins are part of the spliceosome according to the SAFE and GSEA analysis. Using g:Profiler, these proteins were allocated to a certain functional complex. Seven proteins form the CPSF complex and 6 proteins are subunits of the U2 snRNP complex, containing the SF3b complex (SF3B1, SF3B3, SF3B6).

## Notes

### Competing Interest Statement

The authors have declared no competing interest.

